# Comprehensive Cross-Domain Taxonomic Classification of Microbiotas using Partitioned Amplification Multiplexed Amplicon Sequencing (PAMA-seq)

**DOI:** 10.1101/2025.06.18.660470

**Authors:** Xiangpeng Li, Kai R. Trepka, Fangchao Song, Than S. Kyaw, Peter J. Turnbaugh, Adam R. Abate

## Abstract

Microbial communities encompass diverse bacteria, archaea, and eukaryotes that play vital roles in ecosystems and host health. Comprehensive analysis of these communities requires accurate, quantitative, and cross-domain profiling, yet current sequencing methods—metagenome shotgun sequencing (MGS) and ribosomal RNA (rRNA) amplicon sequencing—struggle to achieve these goals in a single assay. MGS provides broad functional insights but suffers from high cost, computational burden, and reliance on incomplete reference databases. rRNA amplicon sequencing, while more taxonomically targeted, typically profiles either prokaryotes or eukaryotes separately, depending on the chosen primer sets, and thus lacks simultaneous cross-domain resolution. To address these limitations, we developed PAMA-seq, a droplet-digital multiplex PCR technique. PAMA-seq partitions DNA sample from microbial communities into nanoliter droplets, each containing a single template molecule, and independently amplifies both the 16*S* and 18*S* rRNA genes, resulting in uniform amplification efficiency and accurate quantification. We validated PAMA-seq on a synthetic microbial community, a stool sample from a patient with colorectal cancer, and a coastal seawater sample. The method provided stable, cross-domain taxonomic profiles at sequencing depths as low as 10^4^ reads—one to two orders of magnitude fewer than comparable shotgun metagenomics approaches—while significantly reducing variability between replicates. By combining comprehensive domain coverage with low sequencing depth requirements, PAMA-seq offers an efficient, cost-effective, and scalable method for monitoring microbial communities across diverse ecosystems, from clinical samples to global marine environments.

## Introduction

Microbial communities comprise diverse and interacting populations of prokaryotic and eukaryotic organisms that collectively shape ecosystem stability and host health^1,2^. Because most microorganisms remain difficult to culture^3,4^, cultivation-free sequencing methods are indispensable for identifying which species are present and quantifying their relative abundances. Comprehensive ecological and clinical insights require simultaneous measurement across all microbial domains—bacteria, archaea, and microbial eukaryotes—yet current sequencing methods rarely achieve accurate, unbiased quantification across domains in a single assay.

Two sequencing approaches currently dominate microbiome profiling: metagenome shotgun sequencing (MGS) and ribosomal RNA (rRNA) amplicon sequencing. MGS sequencing randomly samples genomic DNA from the entire microbial community, revealing extensive information about gene content and metabolic potential^5^. Taxonomical classification of these sequences typically employs DNA-to-DNA, DNA-to-protein, or marker-gene methods^6–9^. While powerful, MGS-based taxonomic classification accuracy strongly depends on large and rapidly expanding reference databases, now containing billions of sequences and trillions of nucleotides^10^. The scale of these databases leads to increased computational complexity, higher false-positive rates^10,11^, and difficulties in classifying organisms absent from databases without extensive computational assembly.

In contrast, rRNA amplicon sequencing specifically targets highly conserved small-subunit genes—16*S* rRNA for bacteria and archaea, and 18*S* rRNA or ITS regions for eukaryotes^12–17^. Due to their well-characterized evolutionary relationships, rRNA sequences can accurately position unknown microbes within existing taxonomic frameworks. Curated rRNA databases, such as SILVA, RDP and Greengenes^18–22^, enhance reliability by systematically removing problematic sequences and ensuring consistent genus-level annotations. However, traditional rRNA amplicon methods usually achieve only genus-level resolution, with species-level precision limited to certain well-characterized groups^23–25^. Furthermore, quantitative accuracy requires corrections for variability in rRNA gene copy numbers (ranging from 1–4 copies in archaea, 1–15 in bacteria and over a hundred in eukaryotes^23,24^). Additionally, attempts to simultaneously amplify prokaryotic (16*S*) and eukaryotic (18*S*) rRNA targets using conventional multiplex PCR face inherent technical challenges^26^. Variations in primer efficiency, GC-content, and amplicon length lead to uneven amplification and selective dropout, disproportionately impacting the detection of rare and low-abundance organisms. These issues severely compromise the reliability of existing cross-domain rRNA sequencing workflows. Thus, a method capable of unbiased, quantitative profiling of microbial communities across domains in a single assay would represent a major advancement for microbial ecology and host-associated microbiome research.

To address these limitations, we developed Partitioned Amplification Multiplex Amplicon Sequencing (PAMA-seq), a droplet-based digital PCR approach that simultaneously quantifies prokaryotic (16*S*) and eukaryotic (18*S*) targets. PAMA-seq partitions microbial DNA into thousands of nanoliter droplets, each containing at most one rRNA gene template. Within each droplet, multiplexed primers independently amplify their isolated template to saturation, thereby eliminating competitive amplification biases and preserving rare sequences. Because each droplet-positive molecule is amplified individually, resulting amplicon counts accurately reflect initial template abundance within each primer set. After normalization for known rRNA gene copy-number variation, PAMA-seq provides precise quantitative community profiles across microbial domains at approximately ten-to hundred-fold lower sequencing depth compared to MGS, achieving practical cost-effectiveness comparable to conventional amplicon sequencing. We demonstrate the broad applicability, sensitivity, and accuracy of PAMA-seq with two distinct microbiome samples: one from a colorectal cancer patient’s gut and another from a coastal seawater environment. These examples highlight PAMA-seq’s robust capacity to sensitively detect and quantify bacteria, archaea, and microbial eukaryotes—including rare or previously unidentified taxa—in a single, efficient assay.

## Results and Discussion

### PAMA-seq Workflow

Minimizing PCR bias in rRNA sequencing is essential for accurate microbial profiling. PAMA-seq tackles this by combining droplet digital PCR with two primer sets targeting prokaryotic 16*S* and eukaryotic 18*S* rRNA genes (**Fig. 1A**). Genomic DNA is diluted and partitioned into millions of nanoliter droplets (Poisson loading, λ ≈ 0.3), such that positive droplets contain a single template molecule—ideal for precise quantification (**Fig. S1**). Inside each droplet, the template is amplified to saturation, resulting in a consistent, endpoint-level amplicon yield independent of exponential-phase kinetics. Provided droplets are of uniform volume, the number of positive droplets—and thus total amplicons—reflects the initial rRNA copy number for each primer set. After pooling, the amplicons undergo standard Illumina library preparation (end repair/A-tailing, adapter ligation, brief PCR; **Fig. S2A**). Because Illumina clustering efficiency depends on fragment length, very short or long inserts can bias representation. To minimize this, the 16*S* and 18*S* amplicons were designed to have similar lengths—445 bp and 449 bp, respectively—ensuring that relative read counts remain proportional across domains. PAMA-seq thus converts droplet counts into accurate measurements of microbial abundance across bacteria, archaea, and eukaryotes, without competitive PCR dropout or length-induced sequencing bias.

**Figure 1:**
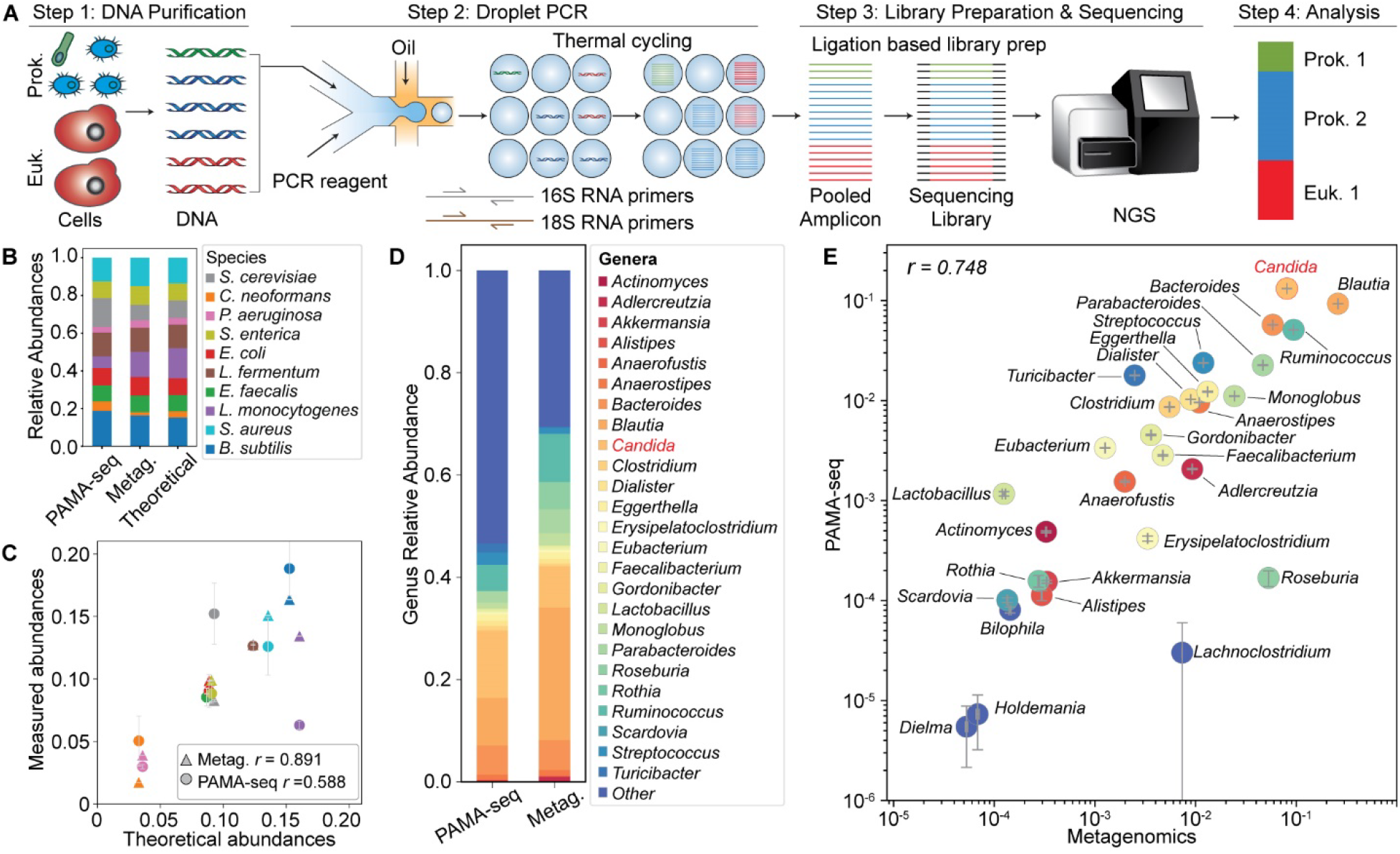
PAMA-seq workflow and validation. (A) Schematic of the PAMA-seq protocol. Genomic DNA is partitioned into nanoliter droplets and amplified via droplet PCR using prokaryotic 16*S* and eukaryotic 18*S* primers (see **Fig. S1**). Sufficient partitioning ensures most droplets contain at most one rRNA template. Amplification reaches saturation, making the number of positive droplets proportional to the initial rRNA copy number. Amplicons are pooled, prepared using standard Illumina library methods (**Fig. S2**), and sequenced—enabling accurate cross-domain taxonomic quantification. (B) Relative species abundances in the ZymoBIOMICS mock community measured by PAMA-seq, metagenomics, and theoretical values. Metagenomic data were normalized for genome size and rRNA-gene copy number. (C) Scatter plot showing Pearson correlation (r) between PAMA-seq, metagenomics, and theoretical abundances. Error bars represent standard deviation from three PAMA-seq replicates and three metagenomic subsamples (2M reads). Analysis of a stool sample from a colorectal cancer patient: (D) Genus-level relative abundances after rRNA copy number normalization determined by PAMA-seq (QIIME 2^28^) and MGS (MetaPhlAn 4^9^), (E) Corresponding correlation plot (r).The eukaryotic genus, *Candida*, is shown in red. Error bars show standard deviation across five metagenomic subsamples (10M reads) and PAMA-seq replicates.

### Validation of PAMA-seq for Quantifying Relative Abundances Across Domains

To benchmark the quantitative accuracy of PAMA-seq, we first tested it on the ZymoBIOMICS Microbial Community Standard, which contains eight bacterial and two yeast species at known proportions. PAMA-seq reads from the ZymoBIOMICS samples were aligned to the reference rRNA gene sequence database. Relative abundances were estimated by normalizing the read counts for each species based on its known rRNA gene copy number (**Fig. 1B, 1C**; **Table S2**). As a comparison, we performed MGS analysis on the same DNA. Reads were aligned to reference genomes and adjusted for both genome length and rRNA-gene copy number (**Fig. 1C**; **Table S3**). Across the ten species, PAMA-seq estimates correlated moderately with expected rRNA abundances (Pearson *r* = 0.59, n = 10; 95% CI 0.06–0.86), while MGS showed better agreement (Pearson *r* = 0.89; 95% CI 0.64–0.97). Notably, the two eukaryotic species, *S. cerevisiae* and *C. neoformans*, exhibited higher-than-expected abundances in the PAMA-seq data. This discrepancy is likely due to the use of incomplete reference genomes for these yeasts; when genome length is underestimated, normalization can artificially inflate the calculated relative abundance. Despite this limitation, the overall results highlight the ability of PAMA-seq to capture both bacterial and fungal taxa with reasonable accuracy. These findings demonstrate that PAMA-seq can be used for cross-domain taxonomic analysis, providing a viable alternative for quantifying the relative abundances of both prokaryotic and eukaryotic species.

### Colorectal Cancer Gut Microbiome Analysis Using PAMA-seq

Next, we applied PAMA-seq to a colorectal cancer patient’s gut sample with known yeast content^27^. We processed reads using QIIME 2^28^ and SILVA v138/v128 databases^18^ (**Table S4**). Bacterial taxa were normalized with rrnDB (The Ribosomal RNA Database v 5.9) copy numbers^29^, and *Candida* reads used the average of the published 18*S* copy numbers (n = 109)^30,31^. MGS reads were analyzed with MetaPhlAn 4.0^9^ (**Table S5**). Twenty-five genera exceeded 0.01% relative abundance in both methods (**Fig. 1D**), yielding strong correlation between the two methods (r = 0.75, p < 0.0001; **Fig. 1E**). Notably, *Candida* exhibited a ∼1.63-fold higher abundance in PAMA-seq compared to MGS, potentially due to uncertainties in 18*S* rRNA copy number—our estimate of 107 based on reported values^30,31^ may not reflect the true value in this sample. Without applying copy number normalization, *Candida* showed a ∼11.58-fold relative abundance in PAMA-seq compared to MGS, and the overall correlation between methods dropped substantially (r = 0.212, **Fig. S3**). These results underscore the critical importance of copy number normalization for accurate quantitative comparisons between amplicon-based and metagenomic profiling approaches. Despite the potential for read inflation in genome-based methods due to *Candida*’s large ∼15 Mb genome^*11*^, PAMA-seq consistently provided sensitive, balanced, and quantitative cross-domain profiles of the gut microbiome. To further improve taxonomic resolution—particularly for eukaryotic microbes—future implementations of PAMA-seq would benefit from the integration of a comprehensive and curated 18*S* rRNA reference database.

### Coastal water microbiome analysis using PAMA-seq

The accuracy of shotgun metagenomic profiling relies heavily on the quality and breadth of reference databases. Many classification tools—whether DNA-to-DNA, DNA-to-protein, or marker-gene based—come with curated, reduced databases that limit their utility in complex environments^10^. While human gut references are relatively complete, environmental collections, especially for marine eukaryotes, remain spotty. In contrast, rRNA databases like SILVA span Bacteria, Archaea, and Eukarya^18,32^, making PAMA-seq an ideal tool for broad environmental assessment.

To illustrate this, we evaluated a coastal water sample using both approaches. Shotgun metagenomic reads were classified with DNA-to-DNA based tools, Kraken 2^33,34^, GOTTCHA 2^35^, and IDseq/CZID^36^; DNA-to-protein based Kaiju^37^; and marker-gene based MetaPhlAn 4.0^9^. PAMA-seq reads were analyzed via QIIME 2^28^ against SILVA v138 and v128^18,32^ (**Fig. 2A**). Shotgun tools classified only a fraction of reads (∼4% by GOTTCHA 2, 35–37% by Kaiju, and ∼59% by CZID), whereas PAMA-seq annotated 98% of amplicon sequences (**Fig. 2B**; **Tables S6–S12**). This contrast reflects shotgun methods sampling all DNA—including non-taxonomic fragments—while PAMA-seq focuses exclusively on rRNA genes. Consequently, PAMA-seq achieved the highest phylum-level Shannon diversity (2.84 vs. 0.36–2.44 for the shotgun methods) (**Fig. 2C**). Only Kaiju, which performs protein-level classification, detected viral taxa with its default database. Eukaryotic phyla were recovered only by Kaiju (with eukaryote-inclusive database) and CZID, and archaeal taxa appeared primarily in Kaiju and Kraken2 output (**Fig. 2D**; **Table S13**). As expected, PAMA-seq detected Bacteria, Archaea, and Eukarya, although it cannot identify viruses.

**Figure 2:**
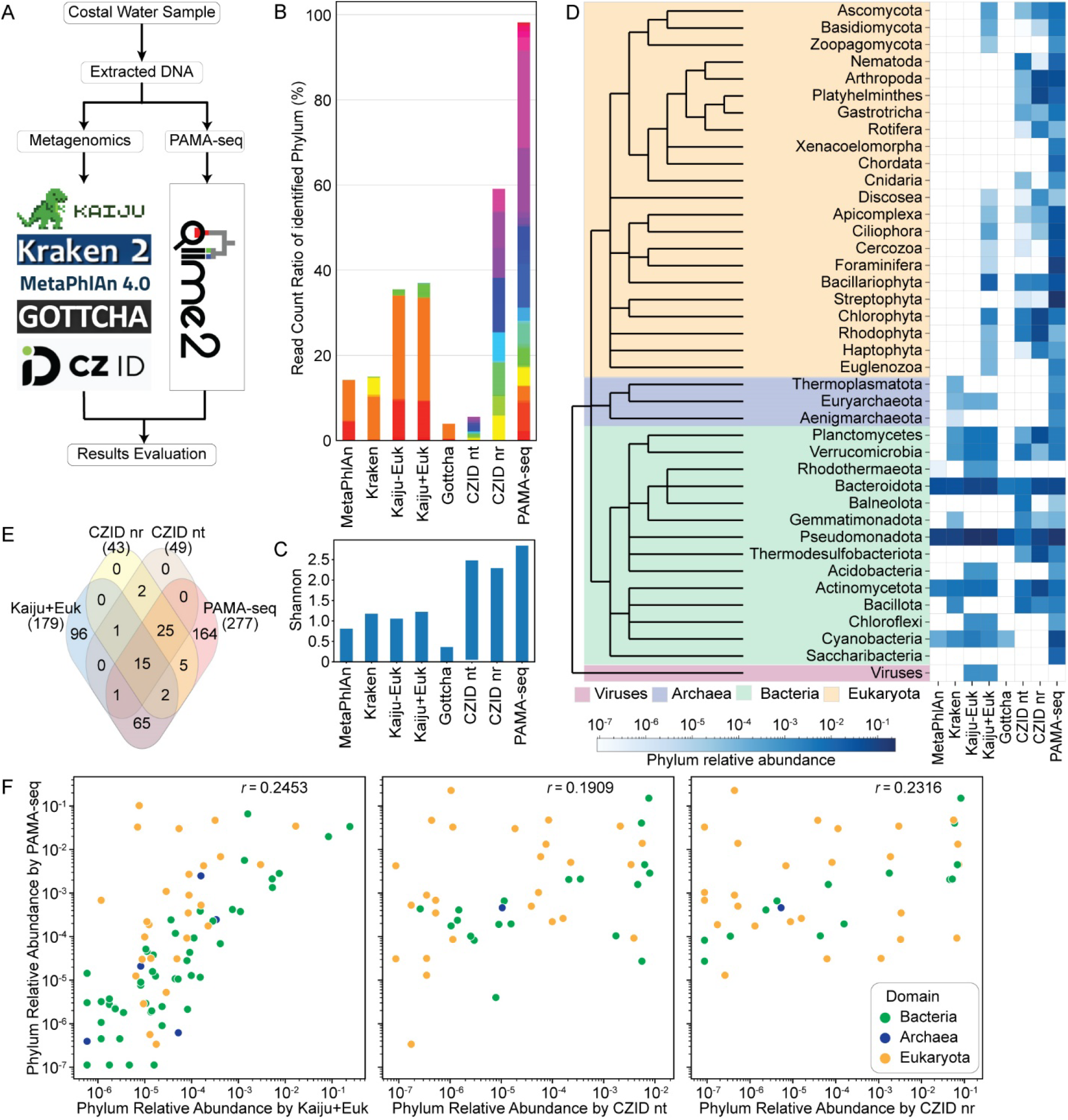
Taxonomic annotation and diversity of a coastal-water microbiome. (A) Study design: DNA from the same coastal-water sample was profiled by shotgun metagenomics and by PAMA-seq. Shotgun reads were classified with Kraken2^33,34^, GOTTCHA2^35^, IDseq/CZID^36^, Kaiju^37^ (NR database with and without eukaryotes), and MetaPhlAn 4.0^9^, while PAMA-seq reads were assigned in QIIME 2^28^ using SILVA v128^18,32^. (B) Read count ratio of identified phylum of the phylum identified in different methods (full legend in **Fig. S4**). (C) Phylum-level Shannon diversity computed for the seven shotgun pipelines and PAMA-seq. (D) Heat-map of phylum abundances (top 40 only) aligned to an NCBI phylogeny; columns, left-to-right: MetaPhlAn 4, Kraken2, Kaiju (NR – euks), Kaiju (NR + euks), GOTTCHA2, CZID-nt, CZID-nr, and PAMA-seq. (E) Venn diagram of phyla shared by the three shotgun pipelines that report eukaryotes and by PAMA-seq. (F) Pairwise correlations of phylum-level abundances between PAMA-seq and those three shotgun pipelines.

PAMA-seq reported up to 277 phyla compared to 179 from Kaiju and 43–49 from CZID, depending on database (**Fig. 2E**; **Table S13**). Across datasets, pipeline correlations ranged from –0.01 to 0.99 (**Fig. 2F, Fig. S5**), illustrating how reference scope shapes outcomes. Some candidate phyla (e.g., *Omnitrophota, Aminicenantota*) and multicellular groups (*Xenacoelomorpha, Chordata*) are sparsely represented in nt/nr and thus often missed by shotgun tools, yet they are recovered by PAMA-seq thanks to its universal rRNA coverage. These results show that PAMA-seq provides deeper, more equitable taxonomic coverage in environmental samples—especially when reference genomes are limited—by leveraging comprehensive rRNA databases and bypassing shotgun biases.

### PAMA-seq requires less sequencing than metagenomics for cross-domain taxonomic analysis

Because shotgun metagenomics sequences all DNA—including many non-taxonomic fragments—while PAMA-seq targets only rRNA sequences, we hypothesized that PAMA-seq would reach stable cross-domain taxonomic profiles with far fewer reads. To test this, we down sampled both the PAMA-seq and MGS datasets to five read depths (10^3^–10^7^ reads), each with five replicates. Shotgun subsets were analyzed using Kaiju^37^ (nr + eukaryotes) and CZID^36^ (nt, nr) because these tools can identify eukaryotes (**Tables S14–S21**), while PAMA-seq reads were reassigned using QIIME 2 and SILVA v138/v128 (**Table S22**).

At higher taxonomic ranks—such as phylum, class, and order—PAMA-seq generally detected a higher Shannon diversity index than shotgun-based methods. In contrast, shotgun sequencing exhibited greater diversity at lower taxonomic ranks, such as genus and species (**Fig. 3A, S6**). This discrepancy is partly due to the tendency of shotgun profilers to inflate species counts when short reads align ambiguously to multiple closely related genomes. Using simulated single-species sequencing data, we found that MGS taxonomic profiling tools like Kaiju and CZID produced high false-positive rates—especially for eukaryotic samples—with many instances showing 0% of reads correctly assigned to the originating species (**Supplementary Note S1**). Second, the targeted 16S/18S regions might lack the resolution to distinguish some congeneric species, a limitation that could be addressed by longer or multi-locus amplicons.

**Figure 3:**
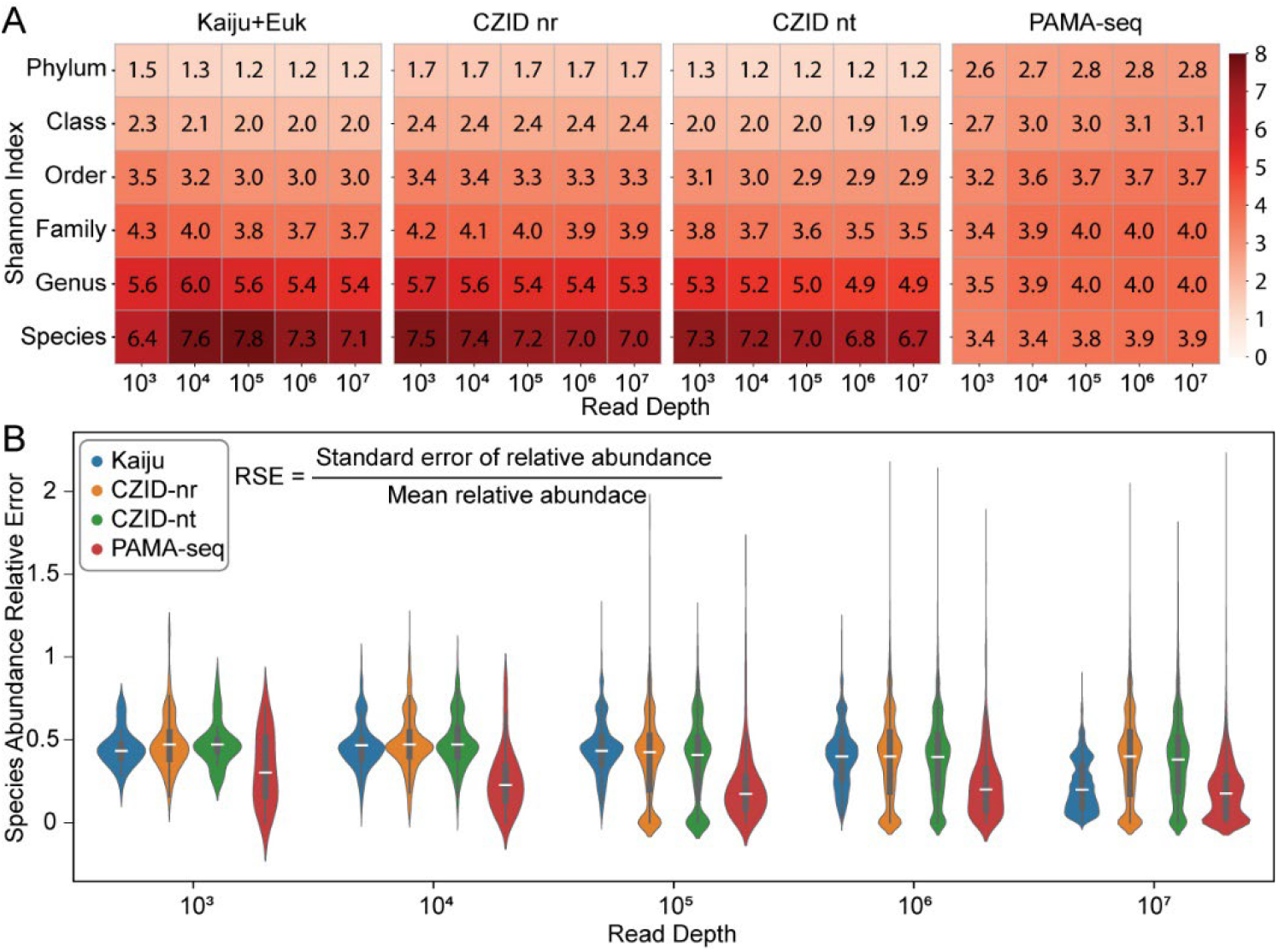
Taxonomic resolution and read-depth effects for PAMA-seq versus shotgun metagenomics. (A) Shannon diversity index each rank (phylum to species) from subsamples of 1 × 10^3^ to 1 × 10^7^ reads (five replicates per depth) using shotgun metagenomics and PAMA-seq. Shotgun data were classified with Kaiju^37^ (nr + eukaryotes) and CZID^36^ (nt, nr); PAMA-seq subsamples were analyzed in QIIME 2. (B) Relative standard error (σ/mean) of species abundances across the five replicates at each depth.

Principal component analysis confirmed that PAMA-seq subsamples clustered more tightly across read depths, indicating greater consistency, whereas shotgun subsamples became increasingly dispersed at lower read counts (**Fig. S6**). We quantified this variability using the relative standard error (RSE = σ/mean), finding that PAMA-seq maintained significantly lower RSE than Kaiju and CZID at all depths, with the greatest advantage below ∼10,000 reads (**Fig. 3B, Fig. S7**). Notably, the RSE of PAMA-seq with 10^4^ reads is significantly lower than that of Kaiju and CZID with 10^5^ or 10^6^ reads across all taxonomic levels. Kaiju with 10^7^ reads could achieved a lower RSE than PAMA-seq with 10^4^ reads (**Fig. 3B, S7**). The high RSE in metagenomics method at low read depth indicates the stochastic sampling effects, biases in low coverage data and analysis methods, which is consistent with usual requirement of more than 10^7^ reads for metagenomic analysis of environmental microbiomes. PAMA-seq achieves comparable cross-domain taxonomic profiles using one to two orders of magnitude fewer reads, offering substantial cost-efficiency for large-scale environmental or clinical surveys.

## Conclusions

We developed PAMA-seq as a sensitive and accurate droplet-PCR tool for cross-domain taxonomic profiling. Across a synthetic community (a simple microbiome with known whole genome sequences of all the members), a colorectal cancer gut microbiome (a complex microbiome with relatively complete reference genomes), and coastal seawater (a complex microbiome with largely incomplete reference genomes), PAMA-seq delivered accurate bacterial, archaeal, and eukaryotic abundance estimates with markedly lower sequencing reads than shotgun metagenomics (10^3^–10^4^ reads versus ≥10^6^), and with far less run-to-run variability, compared to shotgun pipelines. Notably, for microbiomes lacking complete reference genomes, PAMA-seq outperforms all the tested metagenomic methods by detecting all major phyla of bacteria, archaea, and eukaryote. Because each PAMA-seq read is derived from an rRNA gene, nearly all reads contribute taxonomic information, reducing both cost and the variance of diversity metrics. This enables larger, more efficient ecological and clinical surveys using a single streamlined assay. One limitation is that short 16*S* and 18*S* amplicons may not distinguish all closely related species—a drawback that could be addressed through longer or multi-locus primers. Importantly, shotgun metagenomics remains essential for gene-function analysis and virus discovery due to its ability to recover non-rRNA sequences and characterize microbial genomes, making PAMA-seq a powerful, cost-effective complement, rather than a replacement, in multi-domain microbial surveys.

## Experimental

### DNA samples

Synthetic microbial community DNA standard was purchased from Zymo Research (D6305, Lot #213087).

Sea water was collected at Richardson Bay near Sausalito, California (GPS coordinate: 37.8769020, -122.5035899) by submerging a 500 mL sterile bottle into the ocean (March 2024). The sea water was filtered by a 0.45 μm vacuum filter (Millipore, SCHVU01RE) to capture the cells on the membrane. The membrane was cut off from the filter with a sterile razor blade and transferred to a 15 mL centrifuge tube with 5 mL PBS. The cells were released from the membrane by vertexing the tube at maximum speed for 2 min. The cells were then pelleted by centrifuging at 4000 xG for 10 min and resuspended into 500 uL PBS. The DNA was extracted from the cell suspension with QIAamp DNA Stool Mini Kit (Qiagen, 51504) according to the product protocol.

The human fecal sample was collected from a colorectal cancer patient as part of a clinical study (NCT04054908) with full approval by the UCSF Institutional Review Board as described previously^27^. DNA was extracted using the ZymoBIOMICs 96 MagBead DNA Kit. 750 µL of lysis solution was added to a fecal aliquot (50 mg) in a lysis tube. The sample was homogenized with 5 min bead beating (Mini-Beadbeater-96, BioSpec), followed by 5 min room temperature incubation and repeat 5 min bead beating. The sample was centrifuged for 1 min at 15,000 rcf, with 200 µL supernatant transferred into a 1 mL deep-well plate and purified according to the manufacturer’s instructions.

### Multiplex partitioned gene amplification

Commercial or custom-made microfluidics devices can be used to generate droplets for multiplexed PCR. Custom-made microfluidics were fabricated with standard photolithography and soft lithography methods.

400 µL PCR reagents containing 1X Q5® High-Fidelity Master Mix (NEB, M0515), 1% Tween 20, primers (16S-341F: CCTACGGGNGGCWGCAG, 16S-785R: GACTACHVGGGTATCTAATCC, 18S-1183F: AATTTGACTCAACACGGG, 18S-1631R: TACAAAGGGCAGGGACG; 0.1 µM each), and 0.01ng/L DNA sample. were loaded into a 1 mL syringe. The syringe with the PCR reagent and a syringe with 10 mL of droplet generation oil containing 2% PEG-PFPE surfactant (Ran Biotechnologies) in HFE 7500 (3M) were connected to the microfluidic devices. Syringe pumps were used to pump the reagents (PCR reagents at 200 µL/h flow rate and droplet generation oil at 400 µL/h flow rate) into the microfluidic device to form droplets (32 µm in diameter). The droplets are collected into PCR tubes (∼ 50 µL droplets in each tube). The droplet generation oil was removed by pipetting with a gel loading tip and PCR oil containing 5% PEG-PFPE surfactant (Ran Biotechnologies) in FC-40 (3M) was added (50 µL in each tube). The tubes were placed on PCR instrument and thermo-cycled with the following program: 3 min at 95°C for 1 cycle, (15 s at 95°C, 15 s for 55°C, and 1 min at 72°C) for 40 cycles, and 5 min at 72°C for 1 cycle with the lid set at 105°C.

### PAMA-seq library construct and sequencing

The thermal cycled droplets in the PCR tubes were transferred into 1.5 mL centrifuge tubes and the oil layers in the PCR tubes were carefully removed. 20 µL PFO were added and mixed well by vortex to break the emulsion. After centrifuging at 1000 RCF for 1 min, the top aqueous layers in each tube were transferred into new 1.5 mL tubes without disturbing the oil layer and water was added to bring the total volume to 500 µL. The amplicons were purified using 0.7X Ampure XP beads (Beckman Coulter, A63882) and eluted into 50 µL TE buffer. The amplicons were quantified using Qubit™ 1X dsDNA Assay Kits (ThermoFisher, Q33230) and stored at -20°C until the next step.

The sequencing library was constructed using NEBNext® Ultra™ II DNA Library Prep Kit for Illumina® (New England Biolabs #E7645S) according to the product protocol. Briefly, the amplicons were first treated with NEBNext End Prep mix, and the adaptor was ligated. The adaptor-ligated DNA was purified using 0.9X NEBNext Sample Purification Beads. The ligation product was amplified with index PCR (detailed primer sequence in **Table S1**). The indexed sequencing library was purified using 0.7X Ampure XP beads (Beckman Coulter, A63882) and sequenced on Illumina platform.

### Metagenomic sequencing

Metagenomic sequencing data of the ZymoBiomics microbial community DNA standard was provided by Zymo Research (D6305, Lot #213087). The gut microbiome metagenomic sequencing library was prepared using the Illumina DNA Prep Tagmentation Kit (20060059), with quantity and quality checks performed with Quant-iT Picogreen dsDNA Kit (Invitrogen P7589) and TapeStation 4200 (Agilent). The ocean water sample metagenomic sequencing library was prepared using Illumina DNA PCR-Free Library Prep (Tagmentation) kit according to the product protocol. The metagenomic libraries were sequenced using an S1 flow cell on Illumina NovaSeq platform by Novogene (Sacramento, CA).

### PAMA-seq taxonomy abundance estimation

Primers and adapters were removed using the cutadapt trim-paired command in QIIME 2 (v2020.11)^28^. Sequences underwent trimming to 200 bp (forward) or 120 bp (reverse), quality filtering, denoising, and chimera filtering using dada2 (v1.18.0) with QIIME 2 command denoise-paired^38^. Sequence length was filtered to >150 bp with QIIME 2 command feature-table filter-seqs. Taxonomy was assigned to amplicon sequence variants (ASVs) using the SILVA v138 database^39^.

### PAMA-seq compassion with Metagenomics using Zymobiomics sample

The reference genome FASTA files of the ten species of Zymo BIOMICS Microbial Community Standards provided by Zymo Research Corporation (https://s3.amazonaws.com/zymo-files/BioPool/ZymoBIOMICS.STD.refseq.v2.zip). The ten FASTA files of all whole genome references and the ten rRNA gene reference FASTA files were combined into two files and Bowtie2 indices were built using Bowtie2-build command^40^. The metagenomic reads and PAMA-seq reads in FASTQ file were aligned to whole genomes index and rRNA genes index, respectively, using Bowtie2 (v 2.3.5.1) with default setting^40^. The read counts that aligned to each species were calculated using Samtools v1.12 (samtools view) ^41^. The read counts in PAMA-seq were normalized to the copy number of each species’ rRNA copy number and metagenomic-based read counts were normalized to the genome sizes. The relative abundances estimated based on the two methods are then combined for comparison. The relative abundance bar plot and the scatter plot were plotted in Python.

### PAMA-seq compassion with Metagenomics using human gut microbiome sample

For gut microbiome metagenomic data, demultiplexed reads underwent adapter trimming, quality filtering, and merging with FastP (v0.23.2)^42^ and host read removal by mapping to the human genome (GRCh38) with BMTagger (v3.101). Taxonomic abundances were quantified with MetaPhlAn4^9^ using the default settings. For gut microbiome PAMA-seq data, the reads were processed using QIIME 2^28^ as in the previous section.

The PAMA-seq read counts of each taxon were normalized by the rRNA gene copy numbers of the most closely related taxon in rrnDB^29^. In this sample, only one eukaryotic genus was identified, which is *Candida*, and the *Candida* read counts were normalized to the average copy number from previous reports (*C*_*Candida*_ = 106.9091)^30,31^. The relative abundances were then calculated based on the following formula: *p*_*i*_ = (*R*_*i*_/*C*_*i*_)/ ∑_i_(*R*_*i*_/*C*_*i*_), where *p*_*i*_ is the relative abundance for a given taxon, *R*_*i*_ is the read count of a given taxon, *C*_*i*_ is the copy number.

The relative abundances results were then combined and the relative abundance bar plot and the scatter plot were plotted in Python.

### Bioinformatic pipelines evaluation for coastal water microbiome metagenomics

The coastal water metagenomic data was processed either on a local server using MetaPhlAn4^9^ or on web servers, including KBase^43^ (for Kraken2^33^, GOTTCHA2 ^35^, and Kaiju^37^ analysis) and CZID.org (for CZID^36^ analysis).

For MetaPhlAn analysis, demultiplexed reads underwent adapter trimming, quality filtering, and merging with FastP (v0.23.2)^42^. Taxonomic abundances were quantified with MetaPhlAn 4^9^using the default settings.

For the KBase-based analysis, metagenomic paired-end FASTQ files were uploaded to KBase (https://www.kbase.us/)^43^. The reads were first merged using the “Merge Reads Libraries v1.2.2” app. The merged reads were then analyzed using the following apps: GOTTCHA2 v2.1.7 ^35^ (with GOTTCHA viral and bacterial databases), Kaiju v1.9.0^37^ (with NCBI BLAST nr without Euks and NCBI BLAST nr+Euks databases), and Kraken2^33^ (with the Kraken2 database). The results can be accessed at this link (https://narrative.kbase.us/narrative/184052).

For the CZID-based analysis, the paired-end FASTQ files were uploaded and processed using default settings. The analysis results are publicly available (https://czid.org/samples/594122).

The output files from the web servers were downloaded, and relative taxonomic abundances were calculated using the formula: relative abundance of a given taxon = (read counts aligned to that taxon / total read counts). The results from different pipelines were then combined based on the phylum names. The Shannon index were calculated at the species level for each sample using the three methods using the following formular: *H* = −∑_i_[p_i_*ln(p_i_)], where *H* is the Shannon diversity index, and p_i_ is the phylum relative abundance. The read count ration bar plot, the Shannon index bar plot, the heat map of phylum relative abundance, and the phylum level correlation pairplot were plotted in Python using Pandas and Seaborn modules.

The phylogenetic tree was generated and plotted in Python with ETEtoolkit^44^ and Phylo^45^ modules. The Venn diagram was plotted in R using ggVennDiagram library^46,47^ with the list of identified phyla of the four different methods.

### Coastal water microbiome PAMA-seq and metagenomics subsampling analysis

PAMA-seq and metagenomic raw FASTQ files of the costal water microbiome sample were randomly subsampled (1K, 10K, 100K, 1M, and 10M reads per subsample, and 5 sub samples for each read count level) using seqtk (https://github.com/lh3/seqtk). The PAMA-seq subsampled FASTQ files were analyzed as previously described. The coastal water microbiome subsampled FASTQ files were analyzed using Kaiju on Kbase (https://www.kbase.us/)^43^ and CZID (https://czid.org/)^36^. The Kaiju outputs the read counts aligned to different taxon levels in separate files. CZID outputs the read counts aligned to nt and nr database with NCBI ID and the corresponding taxon names at phylum to species levels were identified from NCBI using the NCBI IDs. The resulting taxonomic abundance files using different methods were combined based on taxon names. The identified taxon count from each method were plotted as a heatmap (**Fig. 3A**).

The Shannon index at different taxon levels were calculated at the species level for each sample using the three methods using the following formular: *H* = −∑_i_[*p*_*i*_*ln(*p*_*i*_)], where *H* is the Shannon diversity index, pi is the taxon relative abundance, represents the individual taxa. Scatter plots illustrating the Shannon diversity across different subsamples at various taxonomic levels were generated using Python (**Fig. 3B, Fig. S5**).

Principal coordinate analyses (PCoA) at all taxonomic levels were performed to analyze the beta diversity across different subsamples using Bray-Curtis dissimilarity in Python with scikit-bio module. Scatter plots were generated to visualize the relationships between subsamples and their corresponding read depths. (**Fig. 3C, Fig. S5**).

The relative errors of taxon abundance across different subsamples at the same read depth were calculated using the formula: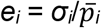, where *e*_*i*_ represents the relative error for the abundance of a specific taxon, *σ*_*i*_ is the standard deviation of the relative abundance across replicate subsamples with the same read depth, and 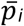 is the mean relative abundance across these replicate subsamples. The distribution of the relative standard errors at different read depth were plotted as a violin plot in Python using Seaborn module (**Fig. 3D, Fig. S6**).

### Simulation for metagenomic accuracy test

The following list of genomic assembly FASTA files were selected and downloaded from NCBI: GCF_000204415.1, GCF_000016525.1, GCF_000007345.1, GCF_000063445.1, GCF_002287195.1, GCF_002310855.1, GCF_000012325.1, GCF_000967895.1, GCF_009362255.1, GCF_001653755.1, GCF_000208865.1, GCF_030704535.1, GCF_000150955.2, GCF_000002765.6, GCF_032362555.1. 150 bp pair-end Illumina simulation reads FASTQ files were generated using ART (https://www.niehs.nih.gov/research/resources/software/biostatistics/art). The read counts for each species are listed in **Table S27**.

All the simulated FASTQ files are processed using Kaiju on Kbase (https://www.kbase.us/)^43^ and CZID (https://czid.org/)^36^, and the outputs (Kaiju, CZID-nt, CZID-nr) were combined as previously stated. The Shannon index were calculated at the species level for each sample using the three methods using the following formular: H = −∑i[pi*ln(pi)], where H is the Shannon diversity index, and pi is the species relative abundance.

## Supporting information

Supplemental Tabels 1-35

## Data availability

All sequencing data is accessible at the NCBI Sequence Read Archive (Accession numbers: PRJNA1232616). Python Jupyter notebooks code used in this paper can be accessed at FSU-Li lab GitHub: (https://github.com/FSU-Li-Lab/PAMA-seq). The supplementary tables can be accessed at FSU-Li lab GitHub and the publisher.

## Author Contributions

X.L. designed the research. X.L., K.R.T., and T.S.K. performed the experiments, X.L., K.R.T., F.S. analyzed the data. X.L. did the data visualization. X.L. and K.R.T. wrote the initial draft of the manuscript. A.R.A. and P.T.J. revised the manuscript. All authors read, reviewed, and approved the manuscript.

## Acknowledgement

This work was supported by the Benioff Center for Microbiome Medicine (BCMM) Trainee Pilot Award (COA7000-138420-7030928-45-A73H5 to X.L.), Mary Anne Koda-Kimble Seed Award for Innovation (MAKK award 2024 to X.L.), and the National Institutes of Health (R01HG008978, U01AI129206, R01AI149699, and R01EB019453 to A.R.A. R01HL122593 and R01CA255116 to P.J.T.). X.L. is supported by Florida State University startup fund. A.R.A. and P.J.T. are Chan Zuckerberg Biohub-San Francisco Investigators. F. S is supported by ENIGMA-Ecosystems and Networks Integrated with Genes and Molecular Assemblies (https://enigma.lbl.gov/), a Scientific Focus Area Program at Lawrence Berkeley National Laboratory, was supported by the Office of Science, Office of Biological and Environmental Research, of the U. S. Department of Energy under Contract No. DE-AC02-05CH1123. The content of this manuscript is solely the responsibility of the authors and does not necessarily represent the official views of the NIH or other funding agency.

## References

1. Tomasulos, A., Simionati, B. & Facchin, S. Microbiome One Health model for a healthy ecosystem. Science in One Health 3, 100065 (2024).

2. Hou, K. et al. Microbiota in health and diseases. Signal Transduct Target Ther 7, 135 (2022).

3. Lagier, J.-C. et al. Current and Past Strategies for Bacterial Culture in Clinical Microbiology. Clin Microbiol Rev 28, 208–236 (2015).

4. Rappé, M.S. & Giovannoni, S. J. The Uncultured Microbial Majority. Annu Rev Microbiol 57, 369–394 (2003).

5. Zhou, J. et al. High-Throughput Metagenomic Technologies for Complex Microbial Community Analysis: Open and Closed Formats. mBio 6, e02288–14 (2015).

6. Camacho, C. et al. BLAST+: architecture and applications. BMC Bioinformatics 10, 421 (2009).

7. NCBI NT database. https://www.ncbi.nlm.nih.gov/nucleotide.

8. McGinnis, S. & Madden, T. L. BLAST: at the core of a powerful and diverse set of sequence analysis tools. Nucleic Acids Res 32, W20–W25 (2004).

9. Blanco-Míguez, A. et al. Extending and improving metagenomic taxonomic profiling with uncharacterized species using MetaPhlAn 4. Nat Biotechnol 41, 1633–1644 (2023).

10. Ye, S. H., Siddle, K. J., Park, D. J. & Sabeti, P. C. Benchmarking Metagenomics Tools for Taxonomic Classification. Cell 178, 779–794 (2019).

11. Sun, Z. et al. Challenges in benchmarking metagenomic profilers. Nat Methods 18, 618– 626 (2021).

12. D’Amore, R. et al. A comprehensive benchmarking study of protocols and sequencing platforms for 16S rRNA community profiling. BMC Genomics 17, 55 (2016).

13. Bradley, I. M., Pinto, A. J. & Guest, J. S. Design and Evaluation of Illumina MiSeq-Compatible, 18S rRNA Gene-Specific Primers for Improved Characterization of Mixed Phototrophic Communities. Appl Environ Microbiol 82, 5878–5891 (2016).

14. Case, R. J. et al. Use of 16S rRNA and rpoB Genes as Molecular Markers for Microbial Ecology Studies. Appl Environ Microbiol 73, 278–288 (2007).

15. Hassler, H. B. et al. Phylogenies of the 16S rRNA gene and its hypervariable regions lack concordance with core genome phylogenies. Microbiome 10, 104 (2022).

16. Stoeck, T. et al. Multiple marker parallel tag environmental DNA sequencing reveals a highly complex eukaryotic community in marine anoxic water. Mol Ecol 19, 21–31 (2010).

17. Bellemain, E. et al. ITS as an environmental DNA barcode for fungi: an in silico approach reveals potential PCR biases. BMC Microbiol 10, 189 (2010).

18. Quast, C. et al. The SILVA ribosomal RNA gene database project: improved data processing and web-based tools. Nucleic Acids Res 41, D590–D596 (2012).

19. Cole, J. R. et al. Ribosomal Database Project: data and tools for high throughput rRNA analysis. Nucleic Acids Res 42, D633–D642 (2014).

20. Guillou, L. et al. The Protist Ribosomal Reference database (PR2): a catalog of unicellular eukaryote Small Sub-Unit rRNA sequences with curated taxonomy. Nucleic Acids Res 41, D597–D604 (2012).

21. O’Leary, N. A. et al. Reference sequence (RefSeq) database at NCBI: current status, taxonomic expansion, and functional annotation. Nucleic Acids Res 44, D733–D745 (2016).

22. DeSantis, T. Z. et al. Greengenes, a Chimera-Checked 16S rRNA Gene Database and Workbench Compatible with ARB. Appl Environ Microbiol 72, 5069–5072 (2006).

23. Langille, M. G. I. et al. Predictive functional profiling of microbial communities using 16S rRNA marker gene sequences. Nat Biotechnol 31, 814–821 (2013).

24. Johnson, J. S. et al. Evaluation of 16S rRNA gene sequencing for species and strain-level microbiome analysis. Nat Commun 10, 5029 (2019).

25. Hugerth, L. W. et al. Systematic Design of 18S rRNA Gene Primers for Determining Eukaryotic Diversity in Microbial Consortia. PLoS One 9, e95567 (2014).

26. Banos, S. et al. A comprehensive fungi-specific 18S rRNA gene sequence primer toolkit suited for diverse research issues and sequencing platforms. BMC Microbiol 18, 190 (2018).

27. Trepka, K. R. et al. Expansion of a bacterial operon during cancer treatment ameliorates fluoropyrimidine toxicity. Sci Transl Med 17, eadq8870 (2025).

28. Bolyen, E. et al. Reproducible, interactive, scalable and extensible microbiome data science using QIIME 2. Nat Biotechnol 37, 852–857 (2019).

29. Stoddard, S. F., Smith, B. J., Hein, R., Roller, B. R. K. & Schmidt, T. M. rrnDB: improved tools for interpreting rRNA gene abundance in bacteria and archaea and a new foundation for future development. Nucleic Acids Res 43, D593–D598 (2015).

30. Rustchenko, E. P., Curran, T. M. & Sherman, F. Variations in the number of ribosomal DNA units in morphological mutants and normal strains of Candida albicans and in normal strains of Saccharomyces cerevisiae. J Bacteriol 175, 7189–7199 (1993).

31. Wickes, B. et al. Physical and genetic mapping of Candida albicans: several genes previously assigned to chromosome 1 map to chromosome R, the rDNA-containing linkage group. Infect Immun 59, 2480–2484 (1991).

32. Yilmaz, P. et al. The SILVA and “All-species Living Tree Project (LTP)” taxonomic frameworks. Nucleic Acids Res 42, D643–D648 (2014).

33. Wood, D. E., Lu, J. & Langmead, B. Improved metagenomic analysis with Kraken 2. Genome Biol 20, 257 (2019).

34. Lu, J. et al. Metagenome analysis using the Kraken software suite. Nat Protoc 17, 2815– 2839 (2022).

35. Freitas, T. A. K., Li, P.-E., Scholz, M. B. & Chain, P. S. G. Accurate read-based metagenome characterization using a hierarchical suite of unique signatures. Nucleic Acids Res 43, e69–e69 (2015).

36. Kalantar, K. L. et al. IDseq—An open source cloud-based pipeline and analysis service for metagenomic pathogen detection and monitoring. Gigascience 9, giaa111 (2020).

37. Menzel, P., Ng, K. L. & Krogh, A. Fast and sensitive taxonomic classification for metagenomics with Kaiju. Nat Commun 7, 11257 (2016).

38. Callahan, B. J. et al. DADA2: High-resolution sample inference from Illumina amplicon data. Nat Methods 13, 581–583 (2016).

39. Yarza, P. et al. Uniting the classification of cultured and uncultured bacteria and archaea using 16S rRNA gene sequences. Nat Rev Microbiol 12, 635–645 (2014).

40. Langmead, B. & Salzberg, S. L. Fast gapped-read alignment with Bowtie 2. Nat Methods 9, 357–359 (2012).

41. Li, H. et al. The Sequence Alignment/Map format and SAMtools. Bioinformatics 25, 2078– 9 (2009).

42. Chen, S., Zhou, Y., Chen, Y. & Gu, J. fastp: an ultra-fast all-in-one FASTQ preprocessor. Bioinformatics 34, i884–i890 (2018).

43. Arkin, A. P. et al. KBase: The United States Department of Energy Systems Biology Knowledgebase. Nat Biotechnol 36, 566–569 (2018).

44. Huerta-Cepas, J., Dopazo, J. & Gabaldón, T. ETE: a python Environment for Tree Exploration. BMC Bioinformatics 11, 24 (2010).

45. Robinson, O., Dylus, D. & Dessimoz, C. Phylo.io : Interactive Viewing and Comparison of Large Phylogenetic Trees on the Web. Mol Biol Evol 33, 2163–2166 (2016).

46. Gao, C.-H., Yu, G. & Cai, P. ggVennDiagram: An Intuitive, Easy-to-Use, and Highly Customizable R Package to Generate Venn Diagram. Front Genet 12, 706907 (2021).

47. Gao, C. et al. ggVennDiagram: Intuitive Venn diagram software extended. iMeta 3, e177 (2024).

