## Supplemental Information for "Comprehensive Cross-Domain Taxonomic Classification of Microbiotas using Partitioned Amplification Multiplexed Amplicon Sequencing (PAMA-seq)"

* Corresponding authors

**Supplementary information** contains:

**Supplementary Notes**

Note S1: Metagenomic taxonomic tool evaluation with single species generated pair end reads.

**Supplementary Figures**

Figure S1: Image of the droplets for PAMA-seq. Scale bar = 100 µm.

Figure S2: Sequencing library preparation procedure.

Figure S3: Taxonomic analysis of a stool sample from a colorectal cancer patient.

Figure S4: Relative abundance of the phylum identified in metagenomics using all different methods and PAMA-seq.

Figure S5: Correlation analysis between the phylum abundance identified using MetaPhlAn, Kraken, Kaiju-euk, Kaiju+euk, Gottcha, CZID-nt, and CZID-nr for metagenomics, and PAMA-seq.

Figure S6: **Alpha Diversity and PCoA Analysis of Metagenomic and PAMA-seq Subsampling at Various Taxonomic Levels.**

Figure S7: Taxonomic abundance estimation relative error across different samples at different read depth.

Figure S8: Strain phylogenetic tree for the species selected for simulation analysis for evaluating metagenomic taxonomic annotation pipelines.

Figure S9: Simulation analysis evaluating metagenomic taxonomic annotation pipelines.

**Supplementary Tables** (S2-S33 are in separated files)

tableS01_primers.csv

tableS02_Zymo_PAMA-seq_aligned_read_counts.csv

tableS03_Zymo_metagenomics_aligned_read_counts.csv

tableS04_Gut_PAMA-seq.csv

tableS05_Gut_metagenomics.csv

tableS06_ocean_meta_metaphlan_output.csv

tableS07_ocean_meta_kraken2_result.csv

tableS08_ocean_meta_kaiju_no_euk.csv

tableS09_ocean_meta_kaiju_with_euk.csv

tableS10_ocean_meta_Gottcha.csv

tableS11_ocean_meta_czid.csv

tableS12_ocean_PAMA-seq_qiime2_result.csv

tableS13_ocean_meta_all_method_and_PAMA-seq_combined_phylum_level.csv

tableS14_ocean_subsample_czi_nt_output.csv

tableS15_ocean_subsample_czi_nr_output.csv

tableS16_ocean_subsample_kaiju_phylum_level_output.csv

tableS17_ocean_subsample_kaiju_class_level_output.csv

tableS18_ocean_subsample_kaiju_order_level_output.csv

tableS19_ocean_subsample_kaiju_family_level_output.csv

tableS20_ocean_subsample_kaiju_genus_level_output.csv

tableS21_ocean_subsample_kaiju_species_level_output.csv

tableS22_ocean_subsample_PAMA-seq_output.csv

tableS23_ocean_subsample_all_method_identified_shannon_index.csv

tableS24_ocean_subsample_all_method_percentage_of_annotated_reads.csv

tableS25_ocean_subsample_all_methods_alpha_diversity.csv

tableS26_ocean_subsample_all_method_relative_error.csv

tableS27_simulation_taxon_info.csv

tableS28_simulation_output_czid_nr.csv

tableS29_simulation_output_czid_nt.csv

tableS30_simulation_output_kaiju_phylum_level.csv

tableS31_simulation_output_kaiju_class_level.csv

tableS32_simulation_output_kaiju_order_level.csv

tableS33_simulation_output_kaiju_family_level.csv

tableS34_simulation_output_kaiju_genus_level.csv

tableS35_simulation_output_kaiju_species_level.csv

**Supplementary Note S1:** Metagenomic taxonomic tool evaluation with single species generated pair end reads.

Given the vast number of reference sequences in metagenomic databases, taxonomic estimation tools are susceptible to false-positive identifications. To assess this issue, we conducted a simulation study using paired-end reads generated from downloaded single-species genomes. Fifteen genomes were randomly selected from NCBI, representing five from each domain: eukaryotes, bacteria, and archaea (**Fig. S8**, **Table S27**). Paired-end reads were simulated using ART (<https://www.niehs.nih.gov/research/resources/software/biostatistics/art>), with read depths ranging from 100,000 to 8,273,014 for each genome (**Table S27**). The simulated samples were analyzed using Kaiju^1^ (NCBI nt database including eukaryotes), CZID^2^ (NCBI nt database), and CZID ^2^ (NCBI nr database) (**Tables S28–S35**).

Not surprisingly, the results showed significant false-positive detection across all three methods (**Fig. S9**). We first analyzed taxon counts across taxonomic levels ranging from phylum to species. Bacterial samples exhibited the lowest false-positive rates, archaea samples demonstrated moderate false-positive rates, while eukaryote samples displayed the highest false-positive rates. For example, from the *Vitis vinifera* sample, Kaiju detected 1,131 different species, and from the *Amyelois transitella* sample, Kaiju detected 617 species.

We further analyzed the Shannon diversity index for each sample analyzed by the three methods, with results ranging from 0.002 to 1.72, none of which align with the single-species input (Shannon index should be zero) (**Fig. S9**). Additionally, we examined the read ratio aligned to the correct taxon, and surprisingly, the majority of samples had 0% reads aligned to the correct species (12 out of 15 for CZID, and 13 out of 14 for Kaiju) (**Fig. S9**). These results suggest that both Kaiju and CZID struggle with accurately identifying species, particularly in eukaryotic samples, leading to significant overestimation at lower taxonomic levels.

Furthermore, these results highlight the challenges faced by current taxonomic estimation methods in accurately distinguishing and quantifying species-level diversity, especially in complex environmental samples with diverse microbial communities. It may be nearly impossible to access the taxonomy abundance ground truth using any sequencing method and bioinformatics analysis alone. We believe that the ground truth lies somewhere between the results from metagenomic and PAMA-seq-based taxonomic estimations, as metagenomics provides significantly overestimated results, while PAMA-seq may underestimate them. Thus, PAMA-seq proves to be highly useful for achieving a more balanced and accurate representation of microbial diversity.

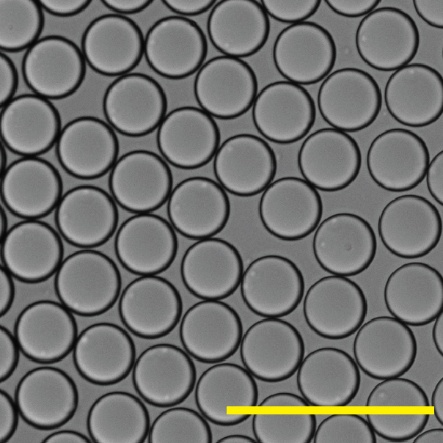

**Figure S1**: Image of the droplets for PAMA-seq. Scale bar = 100 µm.

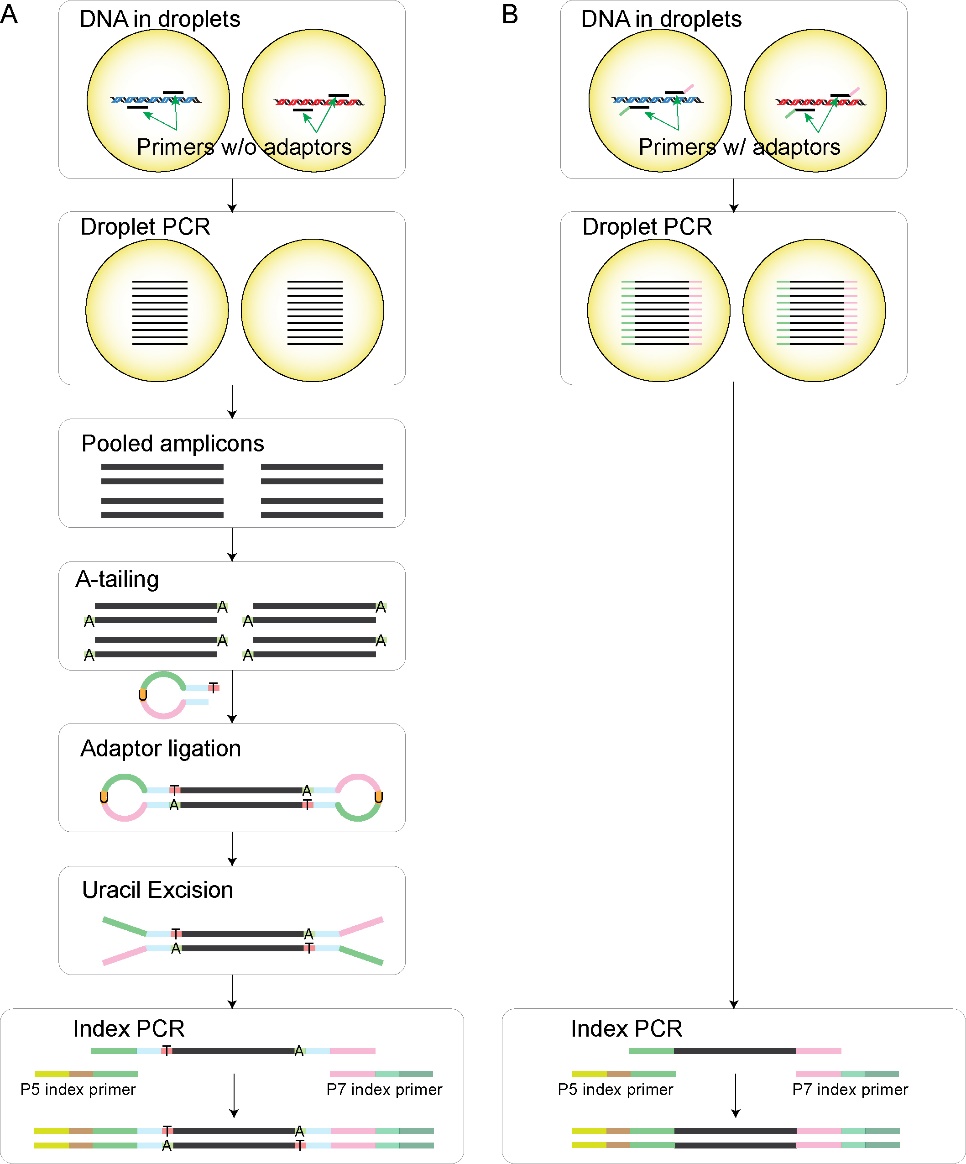

**Figure S2**: Sequencing library preparation procedure. (A) Library preparation method based on ligation. Primers without adaptors are used in the droplet PCR. After pooled all the amplicon together, A-tailing was first introduced to the amplicon molecules. The hairpin adaptor with uracil was then ligated and uracil excision was performed to break the hairpin into the fork adaptor. Index PCR was preformed to generate the sequencing library. (B) Library preparation method based on PCR. Primers with 5’ Illumina adaptors are used in the droplet PCR. After pooling the droplet PCR amplicon, the second PCR was performed to introduce the Illumina P5 and P7 sequencing adaptors. In our preliminary experiments, primer cross-talk was observed when primers shared the same 5' adapters. To avoid this, PAMA-seq uses primers without adapters during droplet PCR (procedure A).

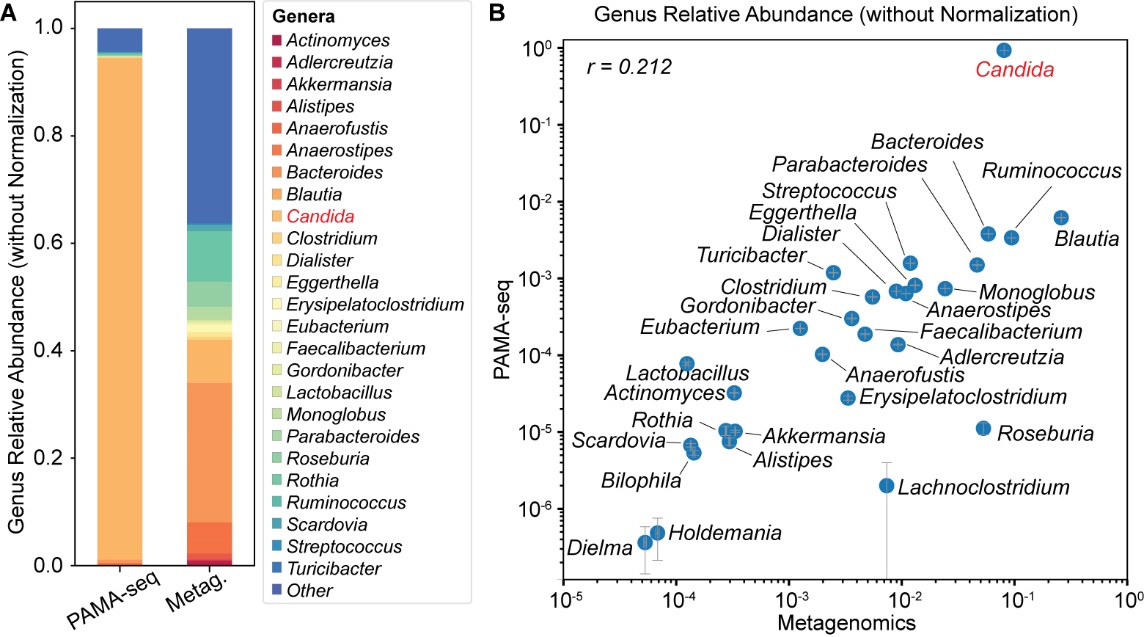

**Figure S3**: Taxonomic analysis of a stool sample from a colorectal cancer patient. (A) Genus-level relative abundances determined by PAMA‑seq and WMS without rRNA gene copy number normalization. (B) Corresponding correlation plot (Pearson r) between the two methods. Error bars represent standard deviation across five metagenomic subsamples (10 million reads each) and technical replicates of PAMA‑seq. Without rRNA copy number normalization, Candida shows disproportionately higher abundance relative to prokaryotic genera.

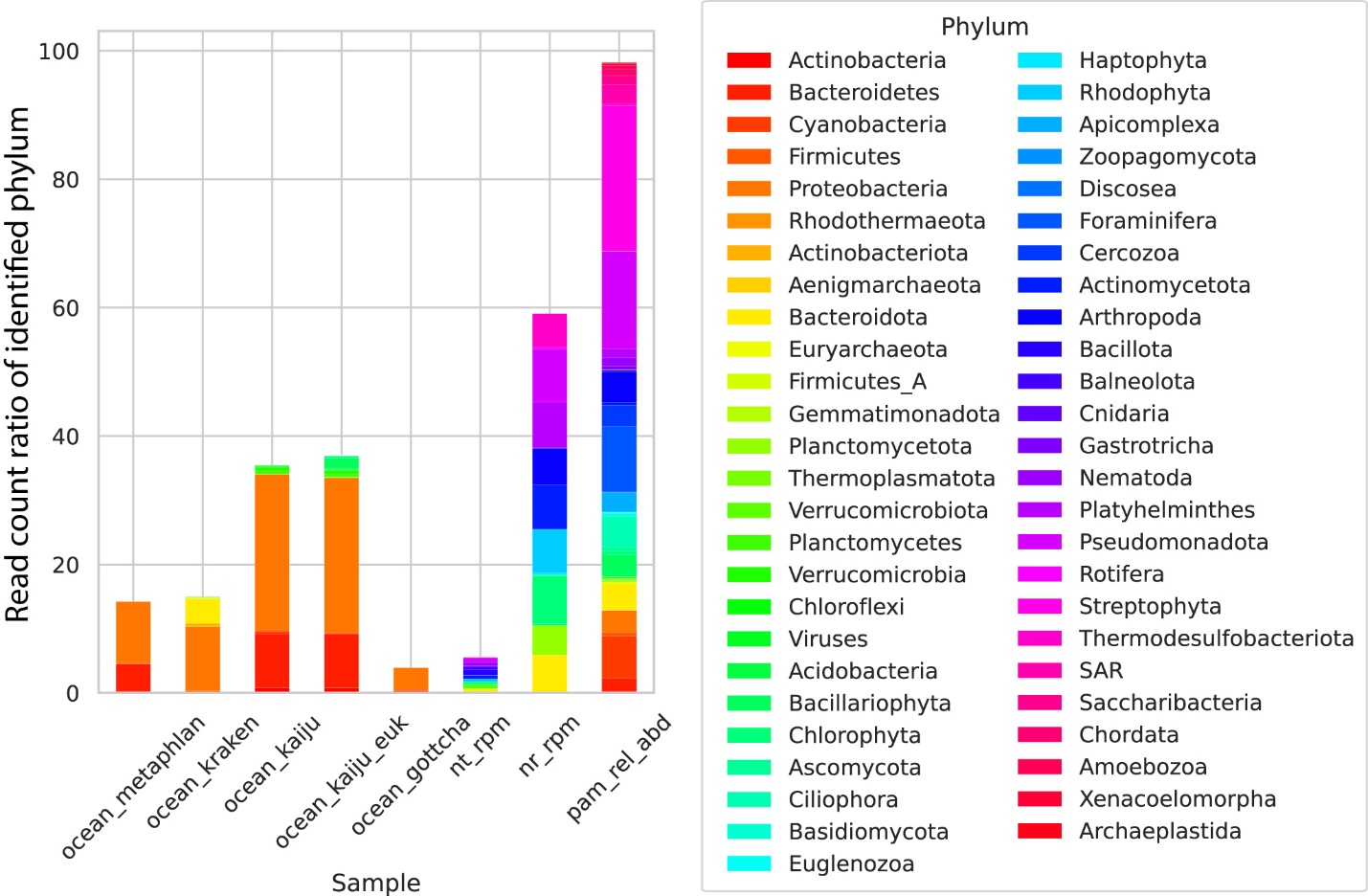

**Figure S4**: Read count ratio of identified phylum of the phylum identified in metagenomics using all different methods and PAMA-seq (same data as **Fig.** **2B**). The legend is included.

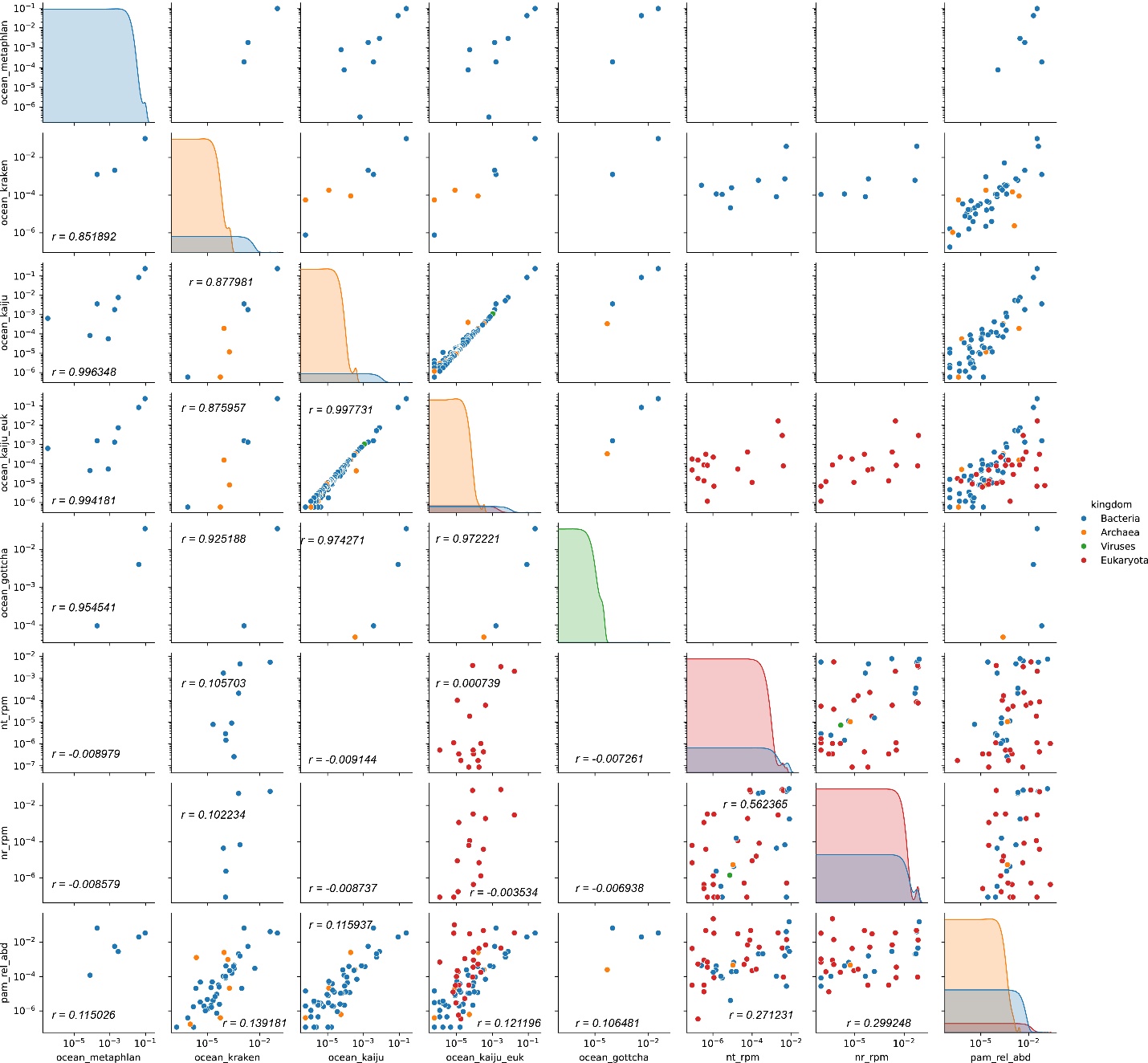

**Figure S5**: Correlation analysis between the phylum abundance identified using MetaPhlAn, Kraken, Kaiju-euk, Kaiju+euk, Gottcha, CZID-nt, and CZID-nr for metagenomics, and PAMA-seq. The dots are color coded by the domain of each phylum.

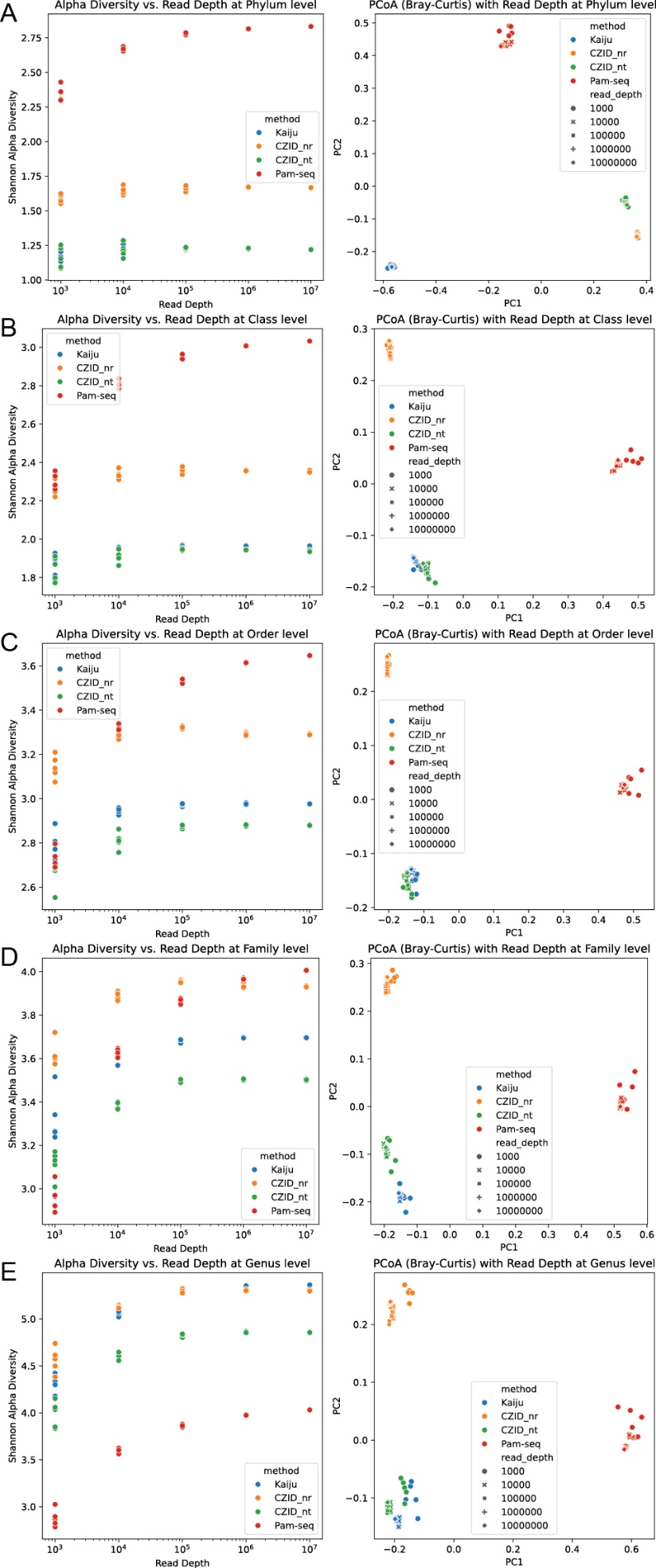
**Figure S6: Alpha Diversity and PCoA Analysis of Metagenomic and PAMA-seq Subsampling at Various Taxonomic Levels.** Subsampling was performed with 1,000, 10,000, 100,000, 1,000,000, and 10,000,000 paired-end reads, with five replicates for each read depth. Three methods were used to analyze the metagenomic subsamples: (1) Kaiju with the nt+euk database, (2) CZID with the nt database, and (3) CZID with the nr database. The Shannon index for each sample at varying read depths and the results of PCoA analysis are displayed across different taxonomic levels: (A) Phylum, (B) Class, (C) Order, (D) Family, and (E) Genus. Analyses at the species level are presented in **Fig. 3B** and **Fig. 3C**.

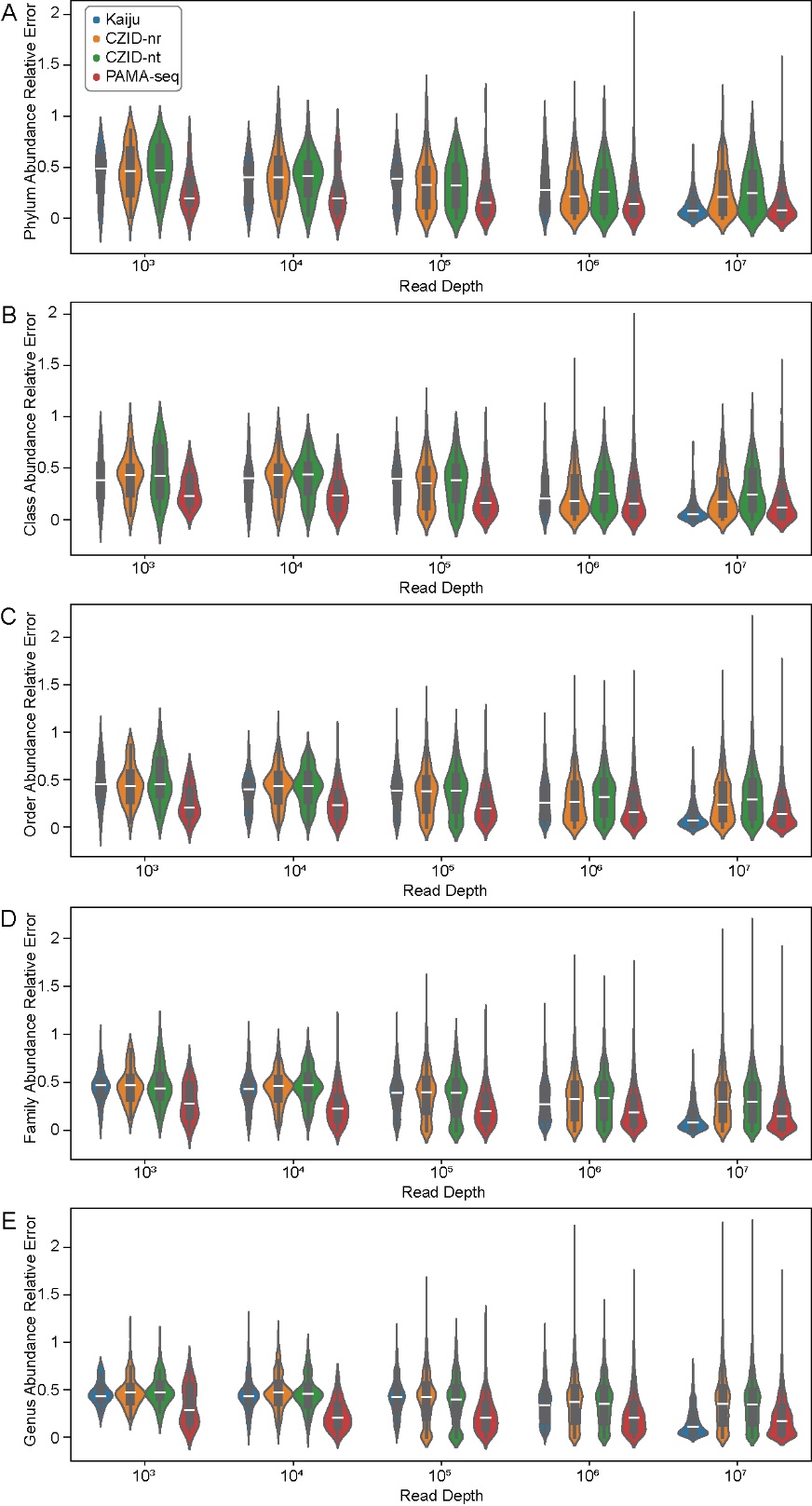

**Figure S7**: Taxonomic abundance estimation relative error across different samples at different read depth. The abundance relative errors are displayed across different taxonomic levels: (A) Phylum, (B) Class, (C) Order, (D) Family, and (E) Genus. Analyses at the species level are presented in **Fig. 3B** and **Fig. 3C**.

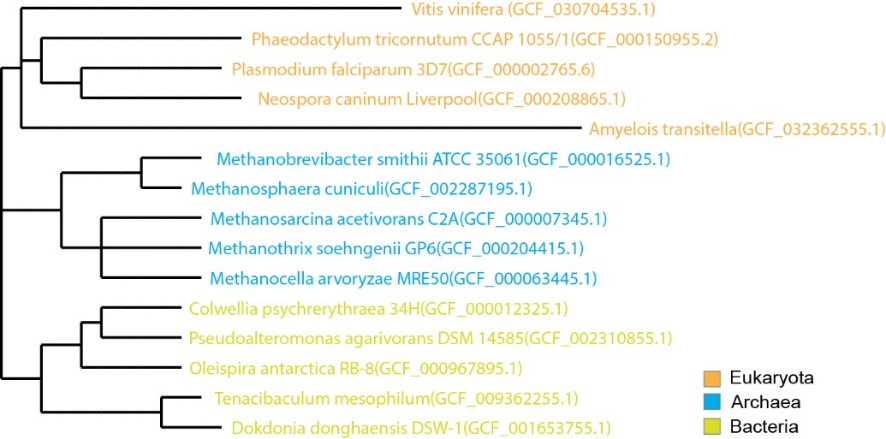

**Figure S8**: Strain phylogenetic tree for the species selected for simulation analysis for evaluating metagenomic taxonomic annotation pipelines. A total of 15 strains were selected for simulation analysis to evaluate the accuracy of metagenomic taxonomic annotation pipelines. Five strains were chosen from each domain: Eukaryote, Archaea, and Bacteria.

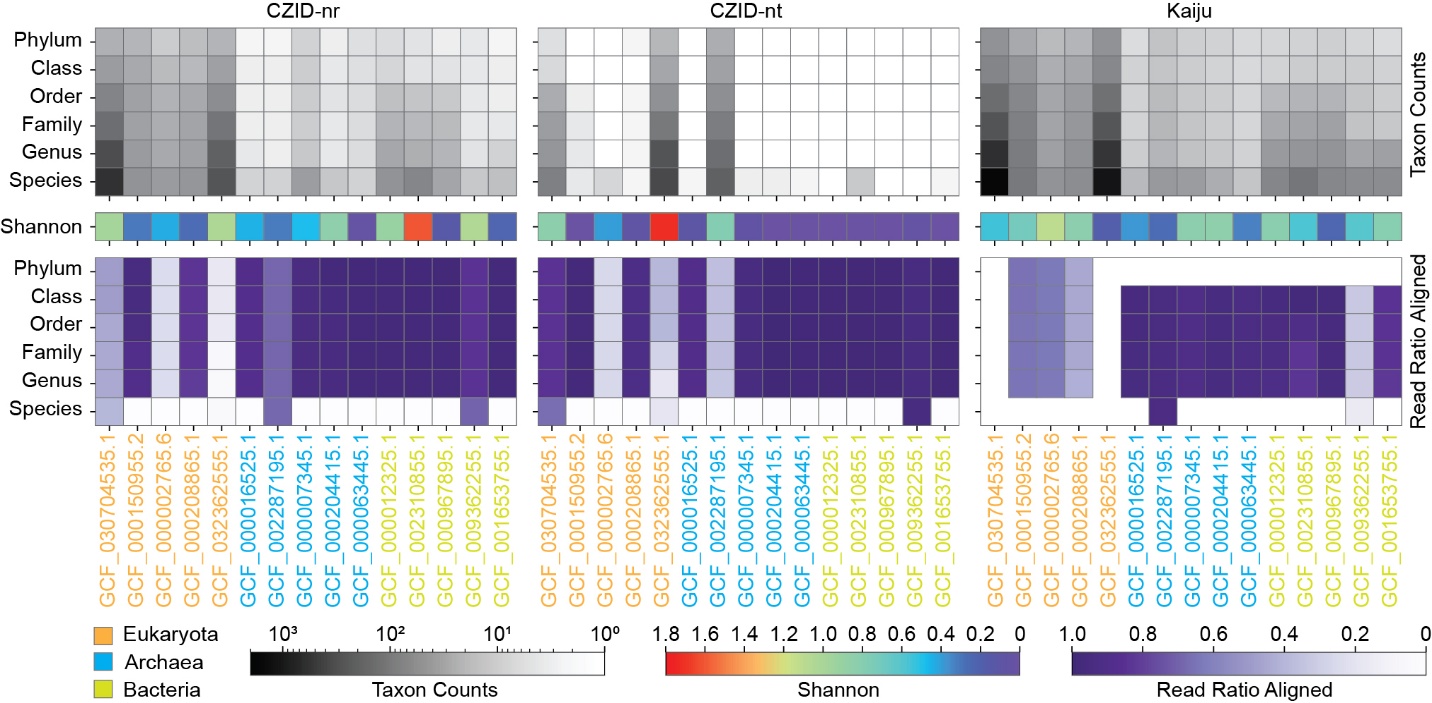

**Figure S9:** Simulation analysis evaluating metagenomic taxonomic annotation pipelines. Heatmaps display (top) the number of identified taxa across different taxonomic levels (gray scale), (middle) Shannon diversity at the species level (rainbow color scale), and (bottom) the proportion of reads used for taxonomic identification (purple color scale) based on simulated reads from 15 species. Left panel: CZID with NCBI-nr database; middle panel: CZID with NCBI-nt database; right panel: Kaiju with default database.

**References:**

1.	Menzel, P., Ng, K. L. & Krogh, A. Fast and sensitive taxonomic classification for metagenomics with Kaiju. *Nat Commun* **7**, 11257 (2016).

2.	Kalantar, K. L. *et al.* IDseq—An open source cloud-based pipeline and analysis service for metagenomic pathogen detection and monitoring. *Gigascience* **9**, giaa111 (2020).
